# Model-based inference of punctuated molecular evolution

**DOI:** 10.1101/852343

**Authors:** Marc Manceau, Julie Marin, Hélène Morlon, Amaury Lambert

## Abstract

In standard models of molecular evolution, DNA sequences evolve through asynchronous substitutions according to Poisson processes with a constant rate (called the molecular clock) or a time-varying rate (relaxed clock). However, DNA sequences can also undergo episodes of fast divergence that will appear as synchronous substitutions affecting several sites simultaneously at the macroevolutionary time scale. Here, we develop a model combining basal, clock-like molecular evolution with episodes of fast divergence called spikes arising at speciation events. Given a multiple sequence alignment and its time-calibrated species phylogeny, our model is able to detect speciation events (including hidden ones) co-occurring with spike events and to estimate the probability and amplitude of these spikes on the phylogeny. We identify the conditions under which spikes can be distinguished from the natural variance of the clock-like component of molecular evolution and from temporal variations of the clock. We apply the method to genes underlying snake venom proteins and identify several spikes at gene-specific locations in the phylogeny. This work should pave the way for analyses relying on whole genomes to inform on modes of species diversification.

## 1 Introduction

Phenotypic variation among organisms is the result of millions of years of genetic evolution (Ho and Zhang, 2018). The evolution of organismal traits is thought to be largely adaptive, even if additional factors such as constraints and historical contingencies (Gould and Lewontin, 1979) or drift for molecular traits (concentrations and molecular functions of non-DNA molecules such as RNAs, proteins, and metabolites) are also recognized (Zhang, 2018). However, whether phenotypic evolution takes place gradually over time or quickly upon speciation is still unclear.

First proposed by paleontologists Eldredge and Gould (1972), the theory of *punctuated equilibrium* has been widely studied and discussed for the evolution of morphological traits. The punctuated equilibrium theory proposes periods of stasis interspersed by large effect transformations, preferentially occurring at speciation events (*cladogenetic changes*). This theory aimed to explain the dramatic morphological changes observed in the fossil record, *i.e.*, the sporadic appearance of new species with substantially different quantitative characters, arising suddenly and remaining nearly unchanged for long periods of time (Stanley 1998), in the light of the allopatric model of speciation of Mayr *et al.* (1954). Under this model of speciation, evolutionary novelties arise in small peripheral isolated populations and are rapidly fixed because of the small population size, explaining the gaps in the fossil record (see Barton and Charlesworth 1984, for a critique of this hypothesis). Conversely, the theory of *phyletic gradualism* supposes that traits evolve in a more continuous manner through small effect mutations arising throughout the lifetime of a species (*anagenetic changes*). This view, inherited from Darwin’s work, postulates that populations gradually differentiate morphologically until they are recognized as separate species (Mayr, 1982).

While gradualism has been the dominant idea in the very beginning of continuous trait evolution modeling (Felsenstein, 1973; Hansen, 1997; Butler and King, 2004), punctualism has recently been considered in modern comparative tools focusing on continuous traits (Bokma, 2002, 2008; Landis *et al.*, 2013; Pennell *et al.*, 2014; Landis and Schraiber, 2017). On the one hand, gradualism is the underlying idea justifying the use of diffusion processes like Brownian motion (BM) or Ornstein-Uhlenbeck (OU) processes to model continuous trait evolution (Pennell *et al.*, 2014). On the other hand, punctualism may be implemented through the use of stochastic processes with jumps, for example Lévy processes (Landis *et al.*, 2013; Landis and Schraiber, 2017).

The opposition between *gradualism* and *punctualism, i.e.*, whether morphological changes occur gradually at the same overall pace or in pulses co-occurring with speciation, can be transposed to molecular evolution. Currently, gradualism seems well anchored as the dominant idea in molecular evolution. The modern literature commonly considers that sequences evolve through the accumulation of isolated substitutions arising as a Poisson process through time, with a rate known as the *molecular clock*, which is supposed to be either constant (a hypothesis called *strict clock hypothesis* by Zuckerkandl and Pauling, 1962), or to vary through time and across lineages (see the review on *relaxed molecular clocks* by Lepage *et al.*, 2007). This family of models relies on an underlying gradualistic view, considering only gradual anagenetic changes, *i.e.*, isolated mutations occurring along gene lineages.

Punctualism has previously been considered as a plausible alternative model of molecular evolution (Webster *et al.*, 2003; Pagel *et al.*, 2006). Starting from the observation that, in trees reconstructed by maximum parsimony, there is often a correlation between the number of substitutions inferred between the root of the tree and a tip and the number of nodes on this path (a phenomenon sometimes called the *node-density artefact*, see Fitch and Beintema, 1990), Webster *et al.* (2003) and Pagel *et al.* (2006) hypothesized that this correlation was due to frequent cladogenetic mutation events. They designed a statistical test aimed at establishing whether this correlation was indeed due to such punctual events or to an artifact in the phylogenetic reconstruction. Unfortunately these first attempts to evaluate the idea of punctuated molecular evolution suffered from methodological artifacts and internal inconsistencies (Brower, 2004; Witt and Brumfield, 2004). Since then, we are not aware of any attempts at detecting or modeling punctual modes of molecular evolution.

Despite the widespread use of models with gradual and anagenetic molecular evolution, punctual evolution occurring at cladogenetic events is expected under two important modes of speciation. (i) Under an ecological mode of speciation, *i.e.*, speciation driven by divergent selection and/or by ecologically-mediated sexual selection, suppressed recombination within genomic islands can enhance the fast fixation of *de novo* mutations, or (ii) under a combinatorial mode of speciation, i.e. speciation driven by the novel assembly of old genetic variants through the introgression of possibly advantageous genetic variation during hybridization with a distant lineage (Box 1 and Fig. 1). In both cases, the burst of substitutions at speciation will appear as an instantaneous spike of molecular evolution at the macro-evolutionary scale. Given increasing empirical evidence for such modes of evolution (Wolf and Ellegren, 2017; Marques *et al.*, 2019), it seems necessary to develop models that account for spikes of molecular evolution at speciation events. In addition, if we are able to detect at which speciation events and on which genes such spikes occur, this will inform us on both modes of speciation and the genes involved, an important goal in the field of speciation genomics (Seehausen *et al.*, 2014).

**Figure 1:**
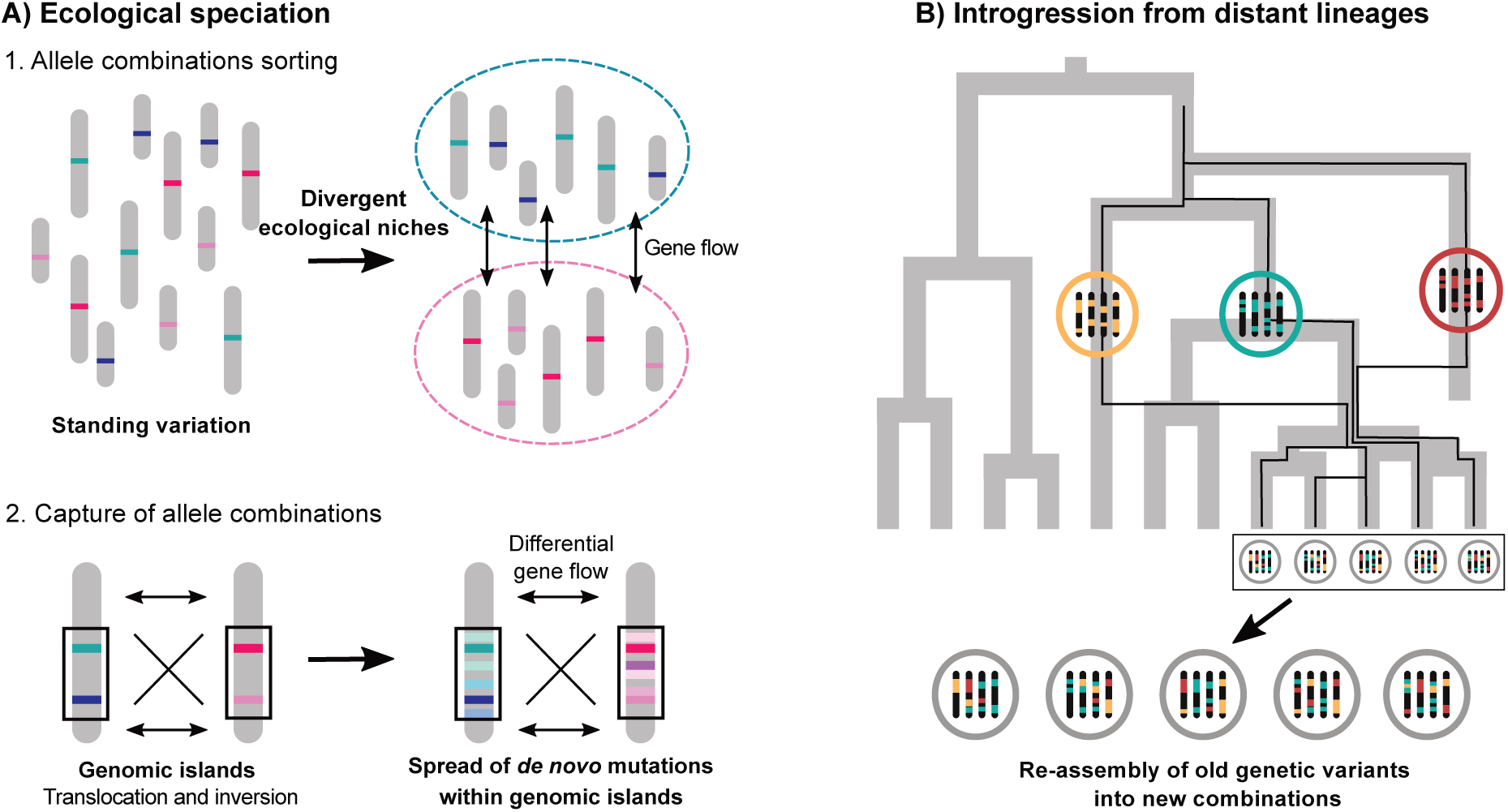
Evolutionary processes causing spikes. (a) Genomic islands formation during ecological speciation with gene flow. Gray sticks represent chromosomes within a population, harboring different alleles (different colors). The dashed ellipses represent two ecologically divergent niches and the arrows indicate gene flow. Genomics islands are delimited by the black rectangles. (b) Introgression from distant lineages. New combinations of alleles between distant lineages lead to rapid speciation and adaptive radiation. Gray tubes represent the species tree, and black lines the gene tree for one sampled gene. Each circle represents an individual from each lineage, and the sticks their genome with a different combination of alleles (different colors). Arrows represent introgression from a lineage with extant and sampled descent (orange) or no sampled descent (red).

### Box 1

**Evolutionary events causing spikes.**

#### 1.1 Ecological speciation with gene flow

Distinct populations of the same species are under strong *divergent selection* when they are specializing to a distinct habitat, to a distinct resource use, or to a different conspecific recognition mechanism, possibly leading to so-called *ecological speciation* (Rundle and Nosil, 2005; Peichel and Marques, 2017).

Before the acquisition of *de novo* mutations, preexisting adaptive variation in quantitative trait loci (QTL) under strong divergent selection can be sorted out through the action of gene flow and selection (Fig. 1a). This mechanism of storing the right alleles in the right habitat requires intermediate levels of gene flow with respect to selection: large enough to perturb local allele frequencies and low enough to avoid homogenization of allele frequencies despite selection acting locally in opposite directions.

Additionally, reduced effective migration close to QTL of larger effect, due to reduced hybrid fitness, will allow close loci of smaller effect to diverge also (divergence hitch-hiking, see Feder and Nosil, 2010). Next, selection can favor factors suppressing recombination between physically distant loci carrying divergently selected alleles. More specifically, QTL can be moved into close genetic linkage through translocations (Yeaman, 2013) and inversions can capture several QTL (Navarro and Barton, 2003; Kirkpatrick and Barton, 2006) (Fig. 1a). In both cases, recombination will be reduced or suppressed in these regions, known as *genomic islands* of speciation, promoting the spread of *de novo* mutations within them.

This rapid formation of highly differentiated genomic islands during ecological speciation (Wolf and Ellegren, 2017) results in an apparent jump of molecular changes localized in the islands (Fig. 1a), *i.e.*, a spike of substitutions. Moreover, genomic divergence will result in the accumulation of between-locus incompatibilities and thus promote hybrid sterility and inviability (Orr, 1997). Therefore, the passive role of differentiation in reproductive isolation guarantees its persistence after speciation is completed.

#### 1.2 Introgression from distant lineages

Long neglected, gene flow (hybridization, horizontal transfer) is now widely recognized between closely related species, and even between distantly related species (Mallet *et al.*, 2016). Gene flow between populations or incipient species, tends to homogenize their allele frequencies, slowing down or preventing speciation. However hybridization between divergent lineages can instead facilitate rapid speciation of few or many (adaptive radiation) lineages (Marques *et al.*, 2019). This process, called *combinatorial mechanism* of diversification, describes the re-assembly of old genetic variants, that have never before been combined together in one population, through adaptive introgression (Fig. 1b). Evidence is accumulating that alleles contributing to reproductive isolation are often much older than actual speciation events, *i.e.*, when populations started to develop reproductive isolation, particularly in cases of rapid speciation and rapid species radiations. Among many other examples (reviewed in Marques *et al.*, 2019), the genomic variation underlying the host switches and associated reproductive isolation of the 200 year-old apple maggot fly *Rhagoletis pomonella* species complex evolved 1.6 million years earlier (Feder *et al.*, 2003; Xie *et al.*, 2008). Similarly, the LWS haplotype polymorphism (in relation to light conditions at different water depths and female mate choice), underlying the cichlid fish radiation in Lake Victoria 100 - 200,000 years ago, emerged by hybridization between two cichlid lineages that diverged about 1.5 million years before (Seehausen *et al.*, 2008; Meier *et al.*, 2017a). Under this scenario of hybridization between two distant lineages, if no descendant of the donor lineage is sampled at present time (introgression from ‘ghost’ lineage), or if gene flow is not taken into account, the introgression event will result in an apparent sudden jump in the molecular evolution of the receiver lineage, *i.e.*, a spike of substitutions.

Here we develop a model of molecular evolution with both gradual mutations and spikes at speciation. In addition, we design a statistical inference protocol in order to infer the parameters of the model and spiking events from molecular sequences associated to a known dated tree. We identify conditions on the values of the model parameters under which spikes can be distinguished from gradual mutations, both in the absence and in the presence of fluctuations of the clock. We evaluate the performance of our inference method on simulated data. Finally, we apply this method to snake venom protein evolution. Venoms, composed of several proteins able to attack biological pathways, are a key adaptation for many snakes that facilitates the capture and the predigestion of prey. Recent evidence suggests that venom composition among snakes evolved in three distinct envenomation strategies, two of them being associated with two distinct families (Barua and Mikheyev, 2019). We test if these two envenomation strategies have left different spiking patterns in associated genes.

## 2 New Approaches

### 2.1 Model of molecular evolution with spikes

Under the idea of punctuated molecular evolution, we develop here a new relaxed clock model built on the joint action of *cladogenetic punctuated evolution* and *anagetic gradual changes*, instead of only anagenetic gradual changes along the tree. We model these episodes of fast accumulation of substitutions as punctual events called *spikes*, which co-occur with speciation events. They are superimposed to standard, strict-clock molecular evolution.

More specifically, we model the diversification of a clade according to a birth-death process with a constant per lineage speciation rate *λ*, constant extinction rate *d* (Fig. 2a) and sampling probability *f* of each extant species. Next, we model the evolution of sequences on the resulting phylogenetic tree for this clade. Sequences are made of *N* homologous nucleotide sites, which evolve both gradually along phylogenetic branches, and punctually at speciation events. Gradually: Along the branches of the phylogeny, the base at each site undergoes a substitution at rate *µ*. Punctually: At each speciation event and for each daughter species, a spike occurs with probability *ν* (Fig. 2a). When a spike occurs, each site experiences a substitution with probability *κ*. Whether gradual or punctual, each substitution is a transition with probability *a* and either of two possible transversions with probability *b* = (1 − *a*)/2.

**Figure 2:**
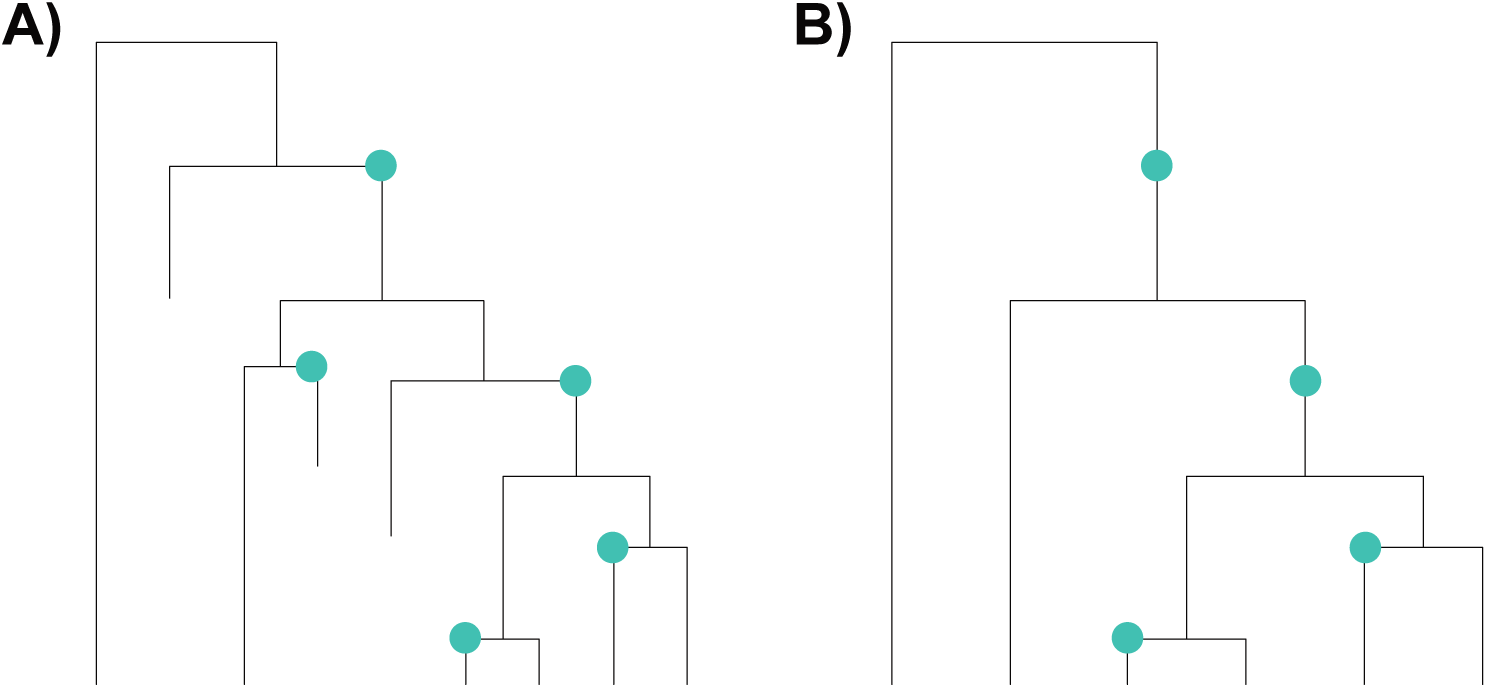
Punctuated model of molecular evolution. Spikes of mutations (green dots) happen at speciation events during the evolution of a clade represented by a phylogenetic tree (A); On the reconstructed phylogeny (B) these spikes occur at nodes, or along branches when one of the two daughter lineages did not leave any sampled descendant.

Since each node in a phylogenetic tree represents a speciation event, our model assumes that a node is associated with a spike when the speciation event has been driven by one of the two processes described in Box 1. This model consists in seeing the genome as a multidimensional trait that may jump at speciation, exactly as in models of quantitative character evolution undergoing ‘shifts’ or ‘jumps’ at speciation (*e.g.* see Bokma, 2008). Seven parameters, described in Box 2, were used to design this relaxed clock model combining a basal molecular evolution with fast accumulation of substitutions (spikes).

#### Box 2

**Model parameters.**

The model of molecular evolution with spikes has seven parameters. They govern the diversification process (*λ, d, f*), basal molecular evolution (*µ* and *a*), the spiking events (*ν*) and substitutions at spikes (*κ*).

- **Speciation rate** (*λ*). Each species independently gives rise to a new species at rate *λ*.
- **Extinction rate** (*d*). Each species independently becomes extinct at rate *d*.
- **Sampling probability** (*f*). Each species extant at present time is independently sampled with probability *f*.
- **Molecular clock** (*µ*). Each base independently undergoes a substitution at rate *µ*.
- **Spike probability** (*ν*). At a speciation event, each of the two daughter lineages undergoes a spike independently with probability *ν*.
- **Substitution probability at a spike** (*κ*). At a spike, each base in the DNA sequence has a probability *κ* to undergo a substitution.
- **Transition and transversion probabilities** (*a* and *b*). Any substitution, gradual or punctual, can be the unique possible transition with probability *a* or either of the two possible transversions with probability *b* = (1 − *a*)/2.

In the remainder of the paper, it will be convenient to reparameterize the model by setting *α* = *aµ* the rate of transitions and *β* = *bµ* the rate of each given transversion, so that 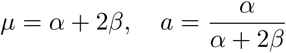 and 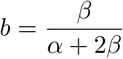.

### 2.2 Distribution of spikes on a reconstructed tree

We call the phylogeny of present-day sampled species 𝒯 (*i.e.*, the reconstructed phylogeny, Fig. 2b). Some spikes can occur along the branches of this reconstructed phylogeny because of hidden speciation events – a speciation event is hidden when it is not seen in the reconstructed phylogeny because one of its two descending clades is extinct or unsampled at present time.

Let 𝒮 denote the number of spikes on each branch of the reconstructed phylogeny 𝒯, including the visible mother node of the branch. We now characterize the law of 𝒮 for a fixed realization of 𝒯.

Time is oriented from the tips (present time *t* = 0) to the root of the tree (time *T*). We call *u*(*t*) the probability that the descent of a species living at time *t* is not sampled at present. This probability is derived in Kendall (1948),

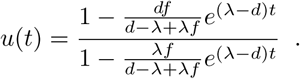

Along any branch of the reconstructed tree, spikes can arise due to hidden speciation events:

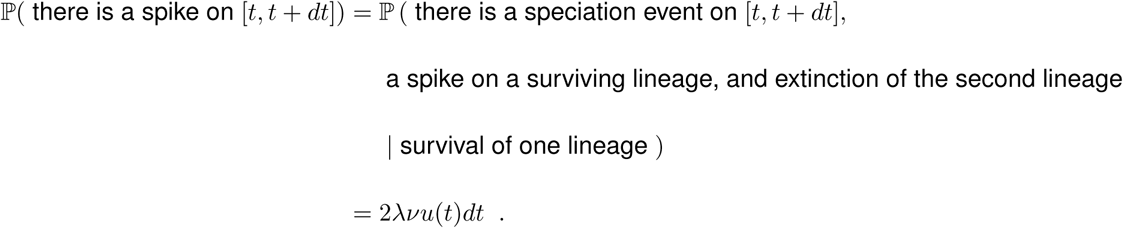

We can thus simulate hidden spikes along a branch of 𝒯 as a Poisson process with rate 2*λνu*(*t*). On a branch originating at time *t*_0_ and ending at time *t*_1_ (with *t*_1_ being closer to the tips than *t*_0_, *t*_1_ *< t*_0_), the number of spikes is Poisson distributed with parameter

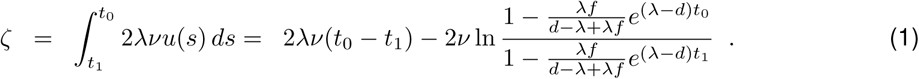

The law of the total number 𝒮 of spikes on each branch of 𝒯 is thus the convolution of a Bernoulli distribution with parameter *ν* (corresponding to the potential spike happening at the visible mother node of the branch) and of a Poisson distribution with parameter *ζ* (corresponding to the spikes occurring at hidden speciation events).

We now need to describe the second ingredient of our model, *i.e.*, the evolution of molecular sequences on a reconstructed spiked tree (𝒯, 𝒮).

### 2.3 Molecular evolution on a reconstructed spiked tree

Conditional on (𝒯, 𝒮), all *N* sites of the sequence evolve independently and identically in distribution. We model basal molecular evolution using the K80 model (Kimura, 1980) unfolding along the tree.

This model is a Markov process with discrete state space {*A, T, C, G*} and rate matrix

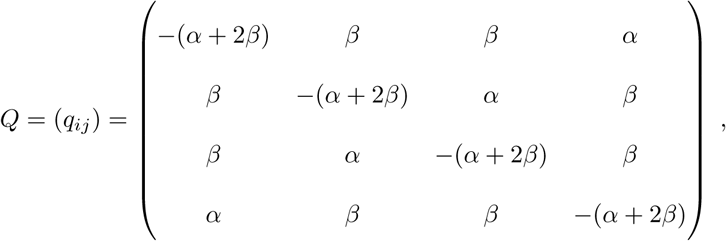

where *α* = *aµ* and *β* = *bµ*. The matrix of transition probabilities between any two nucleotide states separated by time *t* is denoted *P* (*t*) and given by *P* (*t*) = (ℙ(*X*_*t*_ = *j*|*X*_*t*_ = *i*))_*i,j*_ = *e*^*tQ*^.

When a spike occurs, each base mutates with probability *κ* according to the same model of molecular evolution. More precisely, the matrix of the transition probabilities *P*_*S*_ for a nucleotide state just before and just after the spike is defined by

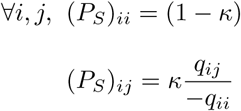

Because *P*_*S*_ and *P* (*t*) commute, the transition probability *P* (*n, t*) of nucleotide states at the extremities of a branch with length *t* and *n* spikes verifies

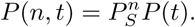

This description of molecular evolution allows us to simulate the evolution of nucleotides along a fixed reconstructed spike tree (𝒯, 𝒮). It also allows us to compute the likelihood of sequences at the leaves, conditional on (𝒯, 𝒮), using a popular *pruning algorithm* (Felsenstein, 1981).

### 2.4 Statistical inference in a Bayesian framework

We consider our model in a Bayesian framework and expose below our strategy to sample from the posterior density. For simplicity, we denote by *p* all probability densities to sketch a Markov Chain Monte Carlo (MCMC) procedure aimed at inferring the joint posterior distribution of parameters and spike positions.

Suppose we fixed the tree realization 𝒯, *i.e.*, we know the tree topology and the times at which branching events occur. We call 𝒜 the alignment of all *N* nucleotides among our *n* extant species. Furthermore, suppose parameters *λ, d, ν, κ, α, β* are not fixed anymore, but are instead drawn from a prior. We fix the following independent uniform priors for these parameters, reflecting the *a priori* knowledge of their range: (i) *λ* and *d* are distributed uniformly on (0, 5), (ii) *ν, κ* are distributed uniformly on (0, 1), (iii) *α, β* are distributed uniformly on (0, 0.1).

The sampling fraction *f* is assumed to be known. We aim at sampling from the *posterior distribution*

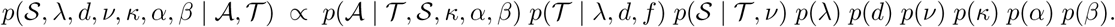

This type of question is classically resolved using a MCMC algorithm to sample the desired distribution. Because we already presented how to compute *p*(𝒮 | 𝒯, *ν*) and *p*(𝒜 | 𝒯, 𝒮, *κ, α, β*), and *p*(𝒯 | *λ, d, f*) is known (Nee *et al.*, 1994), we need only describe two additional components: (i) the initialization of the chain, which provides the first values of the parameters and spike positions, and (ii) the movement proposal, which provides the transitions between two steps of the chain. These are described in Supplementary material (Section A).

We assess visually the convergence of the chain towards its stationary distribution and delete the beginning of the chain (the so-called *burn-in* phase). We use the remainder as an estimate of the target distribution and assess for each parameter the Effective Sample Size (ESS). We implemented the simulation and inference tools in Python, and made this code available in a GitLab repository (https://gitlab.com/MMarc/spike-based-clock/).

### 2.5 Theoretical expectations

Our method aims primarily at distinguishing substitutions arising at spikes from substitutions gradually accumulating between speciation events. Spikes can be statistically indistinguishable from gradual evolution due to two main sources of stochasticity in the process of gradual evolution: variance of the Poisson number of strict clock-like substitutions and heterotachy, that is, temporal variations of the molecular clock itself. These two sources of stochasticity can produce both false negatives and false positives.

The fraction of substitutions associated with spike events (parameter *κ*) has to exceed a certain threshold *κ*_0_ to stay immune from false negatives. If we assume that the clock is relaxed, that is, varies in each branch by a factor *e* which is drawn uniformly in [−*ϵ, ϵ*], then the amplitude of heterotachy *ϵ* has to remain below a certain threshold *ϵ*_0_ to stay immune from false positives.

In Supplementary material (Section B) we propose a generic way of computing the threshold values *κ*_0_ and *ϵ*_0_ for any given parameter set. Namely, Equation (9) yields

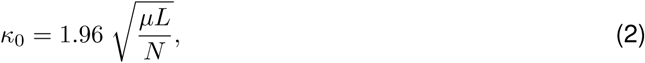

where *L* is the branch length in the phylogeny where a spike occurs, *µ* is the molecular clock and *N* is the sequence length. As a rule of thumb, taking the standard value of *µ* = 10^−2^ per My for the molecular clock, the typical value of *κ*_0_ is

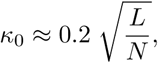

where *L* is measured in My. Conversely, to be able to detect spikes of amplitude 1%, the ratio *L*/*N* must be smaller than 2.5 10^−3^, for example *N* = 1 kb and *L* = 2.5 My, or *N* = 400 bp and *L* = 1 My.

Similarly, Equation (10) in Supplementary material yields for any given *κ*,

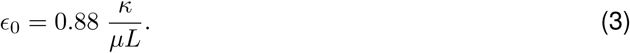

Taking *µ* = 10^−2^, the typical value of *ϵ*_0_ is

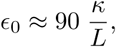

where *L* is measured in My. Conversely, to be able to detect spikes of amplitude 1% in spite of the clock varying by a factor *ϵ* in a branch of length *L*, the product *ϵL* must be smaller than 0.9, for example *ϵ* = 0.3 and *L* = 3 My or *ϵ* = 0.1 and *L* = 9 My.

Instead of artificially letting the molecular clock vary by a fraction ±*ϵ* on each branch, the standard model-based approach of relaxed clock consists of assuming that the clock varies through time like a geometric Brownian motion with infinitesimal variance *σ*^2^. Calculations done in Supplementary material (Section B) yield the following criterion for distinguishing the effect of spikes from that of Brownian-like fluctuating clocks (Equation (12))

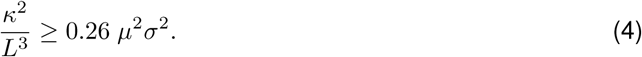

The theoretical predictions displayed in Equations (2), (3) and (4) will be confronted to empirical findings in Section 4.1.

## 3 Results

### 3.1 Performance of the inference method

We checked the ability of the inference method to retrieve parameters under which we simulated artificial datasets (Material & Methods). Parameters of the substitution process (*κ, α, β*) are retrieved quite precisely on each of these simulated datasets (Fig. 3). Parameters (*λ, d*) corresponding to the birth-death process and parameter (*ν*) corresponding to the spike process, have broad posterior distribution, which is expected since each dataset corresponds to only one simulated tree with few branching events (tree size ranged from 12 to 66, with a median of 31.5, Fig. 3). Importantly, the number of spikes, which is our parameter of interest, can be rather well recovered.

**Figure 3:**
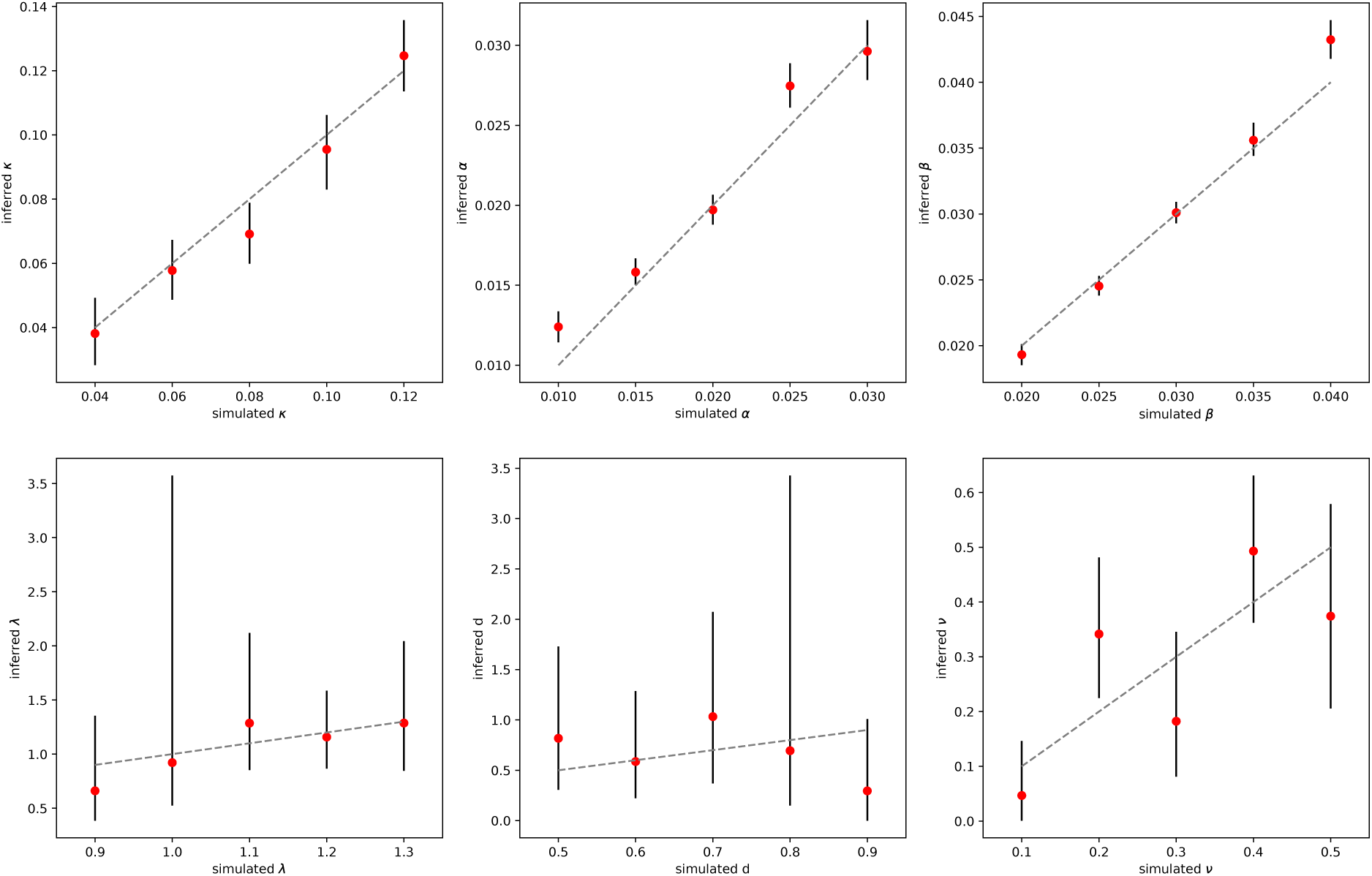
Parameter estimation on a unique dataset simulated under known parameter values. The red dot is the median of the posterior, and the black line represents the 95% envelope of the posterior. Simulated trees in these datasets displayed between 12 and 66 tips and the alignments were 2 kb long.

Our method aims at distinguishing spikes from clock effects. However if the spike amplitude is too low relatively to the stochasticity of the molecular clock, we should not be able to detect spike events (see Section 2.5). We varied *κ* while holding basal substitution rates (*α* = 0.02, *β* = 0.03) constant (Material & Methods). When *κ <* 0.05, most spikes were not detected (Fig. 4a). Conversely, when *κ* > 0.05, all four spikes were correctly inferred (Fig. 4a). For all values of *κ* tested (*κ* ≥ 0.03), no false spike was inferred (Fig. 4b).

**Figure 4:**
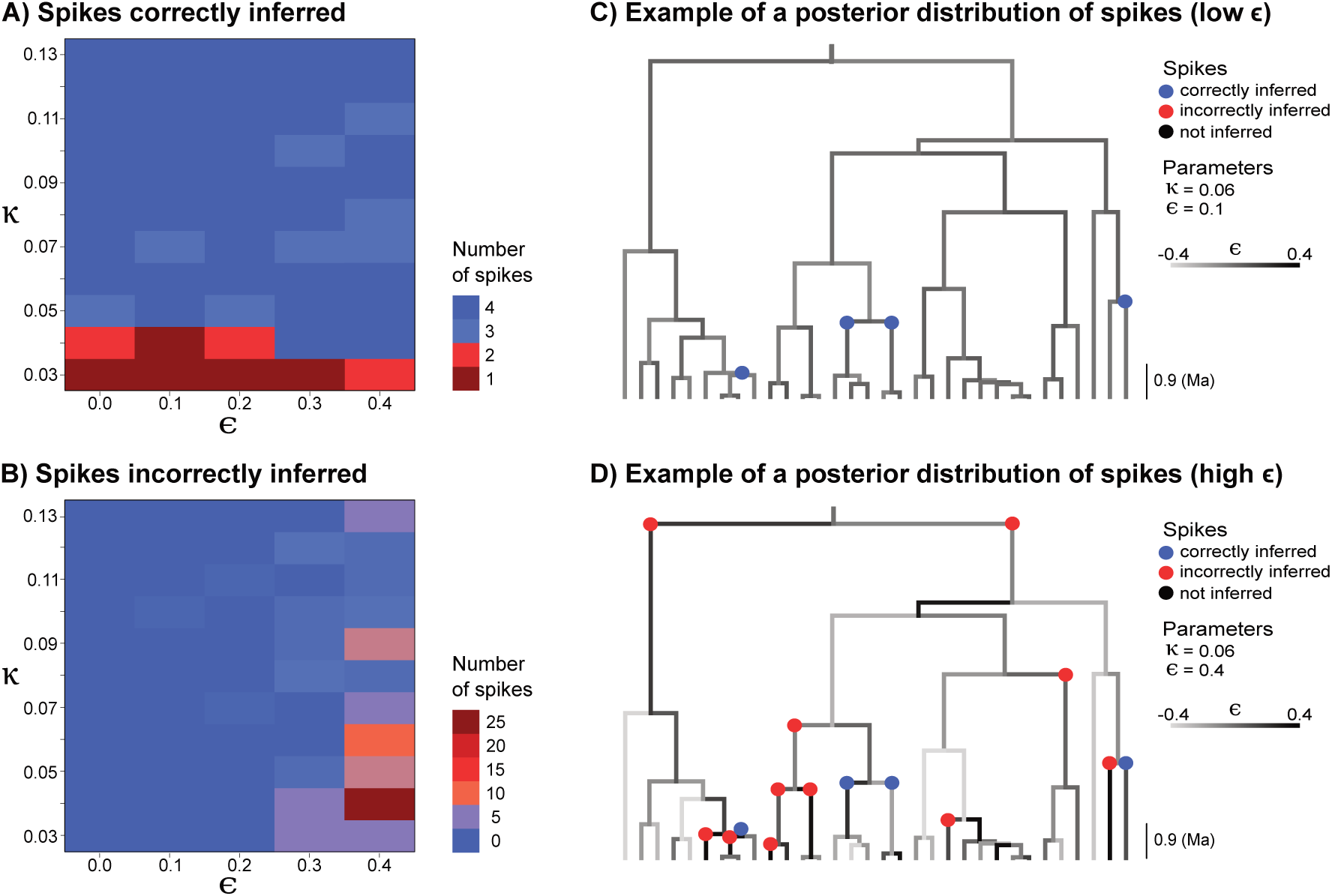
Spike detection accuracy. For a simulated tree with 32 tips and 4 spikes we report (a) the number of these spikes that are detected by the method (true positives) and (b) the number of spikes that are inferred although they did not occur in the simulation (false positives). In (c), the reconstructed tree with spikes for *κ* = 0.06 and *ϵ* = 0.1 (with *κ* the substitution probability at a spike and *ϵ* the amplitude of heterotachy). In (d), the reconstructed tree with spikes for *κ* = 0.06 and *ϵ* = 0.4.

### 3.2 Robustness to model misspecification

Spike amplitude has to exceed some threshold for spike substitutions to be distinguished from clock-like substitutions. This threshold is expected to be larger if the clock itself varies among branches. We tested the sensitivity of the method to a clock varying by a fraction *E* in each branch (Material & Methods). For small departures from a constant molecular clock (*ϵ* ≤ 0.2) and specifically for high substitution probability *κ* at spikes, spikes were correctly inferred (Fig. 4a) with very few false positives (Fig. 4b and c). However, for a large departure from the molecular clock (*ϵ* ≥ 0.3), not all simulated spikes are detected and many false spikes are erroneously inferred (Fig. 4c).

### 3.3 Snake venom proteins evolution

Venom composition among snakes evolved in three distinct envenomation strategies, involving either TFTx (three-finger toxin), SVMP (snake venom mellanoprotease), or the combination of SVSP (snake venom serine protease) and PLA2 (phospholipase A2) (Barua and Mikheyev, 2019). These proteins are not exclusive to snakes. They evolved throughout the Toxicofera clade (Fry *et al.*, 2012) that in addition to snakes includes Anguimorpha (*e.g.*, monitor lizards) and Iguania (*e.g.*, iguanas and chameleons). To test whether these strategies have left differential molecular signatures (spikes) among the Toxicofera, we evaluated the spiking pattern of two proteins: SVSP and CRISP (cystein-rich secretory protein) which like TFTx is a neurotoxin (Yamazaki and Morita, 2004). We selected these two proteins because, unlike SVMP and TFTx, alignments of a substantial number of homologous genes coding for SVSP and CRISP were available that covered a fair part of the Toxicofera clade (Perry *et al.*, 2018). Additionally we analyzed the gene R35 (Orphan G protein-coupled receptor), which is commonly used to reconstruct timetrees as its molecular evolution is near clock-like (Vidal *et al.*, 2007), and so served as a test for false positives.

Within snakes, we found many spikes for CRISP almost exclusively in Elapids and NFFC (non-front fanged colubrids), whereas SVSP spikes were exclusively found in Viperids (see Fig. 5). Interestingly the same pattern was found within Varanids, that is, CRISP spikes are clustered in the clade gathering *Varanus acanthurus, V. gilleni* and *V. scalaris*, and SVSP spikes mainly in the clade gathering *V. gouldii, V. panoptes, V. mertensi* and *V. komodoensis*. Conversely, we did not detect any spike for R35 (supplementary Fig. S1), as expected with this clock-like gene. For virtually each branch, the number of spikes inferred remains very close to an integer after averaging over samples of the MCMC, indicating that the chain does not often switch between equivalent spike configurations but sticks to one optimal configuration throughout the search.

**Figure 5:**
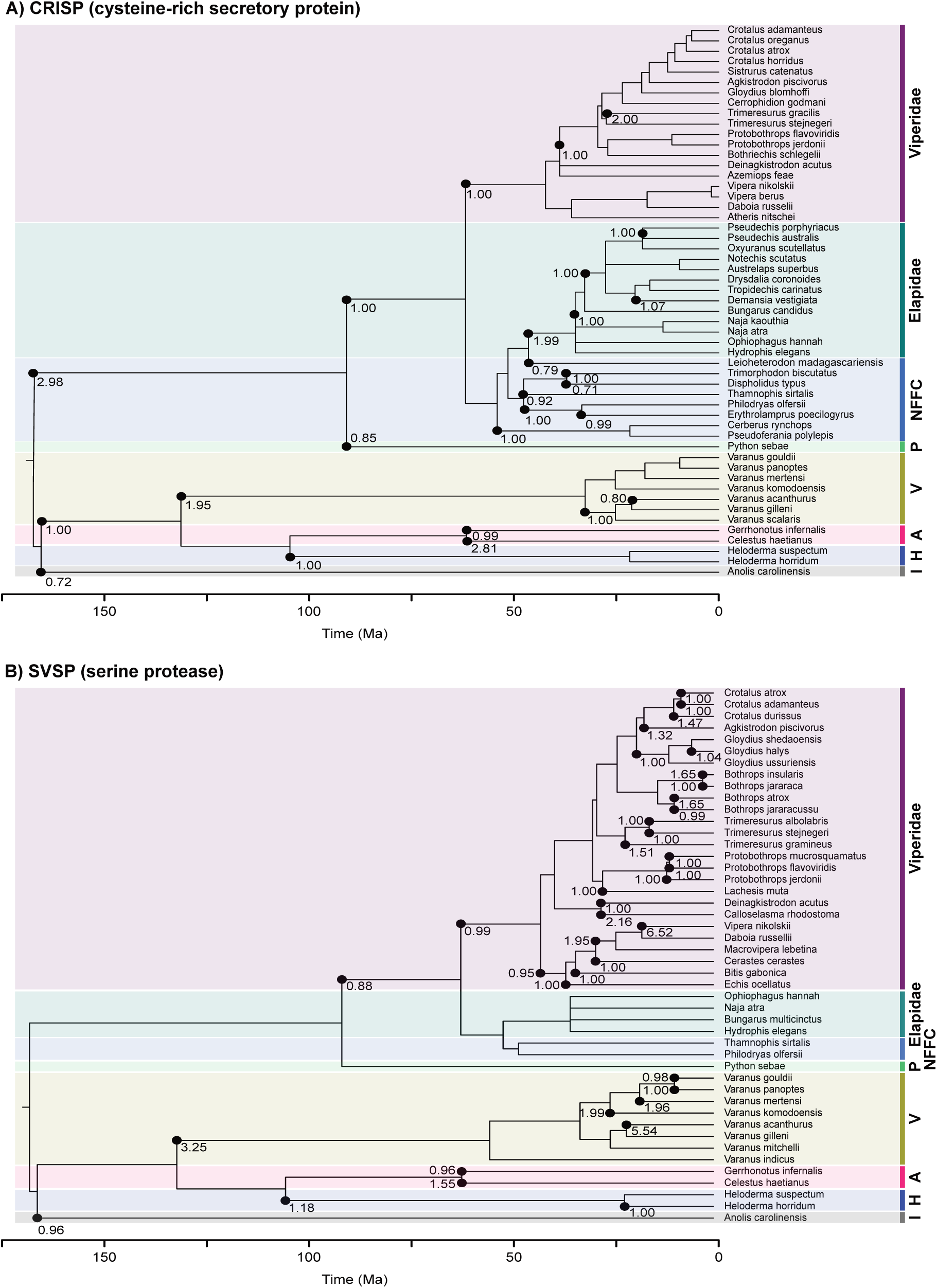
Mutational spikes of snake venom proteins. Spikes were inferred on the venom clade, encompassing among others Iguanidae, Varanidae, and Serpentes, for the CRISP (A) and SVSP (B) proteins. Black dots represent spikes, always placed on their mother node even if they occur along a branch. Numbers indicate the total number of spikes either occurring at the corresponding node or along its descending branch, averaged over the MC chain. NFFC: Non front-franged colubrids, P: Pythonidae, A: Anguidae, H: Helodermatidae, I: Iguanidae.

We found higher substitution rates at the third codon position (parameters *α, β* and *κ*) for SVSP and R35. For CRISP, substitution rates were similar among the three codon positions (see Table 1).

**Table 1:** Parameter values tested. Underlined values correspond to the ones that are fixed when varying the values of another column. For example, we simulated 5 trees and associated alignments with lambda ranging from 0.9 to 1.3 and *d, ν, α, β* and *κ* fixed to the underlined values.

| Parameters | $\lambda$ | $d$ | $\nu$ | $\alpha$ | $\beta$ | $\kappa$ |
| --- | --- | --- | --- | --- | --- | --- |
| Values | 0.9 | 0.5 | <u>0.1</u> | 0.010 | 0.020 | 0.04 |
|  | <u>1.0</u> | 0.6 | 0.2 | 0.015 | 0.025 | 0.06 |
|  | 1.1 | 0.7 | 0.3 | <u>0.020</u> | <u>0.030</u> | <u>0.08</u> |
|  | 1.2 | <u>0.8</u> | 0.4 | 0.025 | 0.035 | 0.10 |
|  | 1.3 | 0.9 | 0.5 | 0.030 | 0.040 | 0.12 |

**Table 2:** Basal substitution rates per codon position (*α*_1_, *α*_2_, *α*_3_, *β*_1_, *β*_2_ and *β*_3_) and substitution probability per codon position at a spike (*κ*_1_, *κ*_2_ and *κ*_3_). We did not report substitution probabilities at a spike for R35 because no spike was detected for this gene.

| Gene | $\alpha_1$ | $\alpha_2$ | $\alpha_3$ | $\beta_1$ | $\beta_2$ | $\beta_3$ | $\kappa_1$ | $\kappa_2$ | $\kappa_3$ |
| --- | --- | --- | --- | --- | --- | --- | --- | --- | --- |
| CRISP | 0.00048 | 0.00044 | 0.00047 | 0.00023 | 0.00021 | 0.00023 | 0.05026 | 0.03503 | 0.05042 |
| SVSP | 0.00039 | 0.00019 | 0.00043 | 0.00020 | 9.473e-05 | 0.00022 | 0.06512 | 0.05911 | 0.05309 |
| R35 | 0.00029 | 0.00019 | 0.00074 | 6.529e-05 | 4.158e-05 | 0.00016 | / | / | / |

## 4 Discussion

### 4.1 Validity and range of application of the method

The model of molecular evolution presented here combines two processes: standard, gradual accumulation of substitutions and spikes occurring at speciation. It is characterized by:

- The basal rate of gradual molecular evolution *µ*, called molecular clock,
- The probability *ν* that a speciation event results in a spike on the sequence under scrutiny,
- The mean amplitude *κ* of spikes, which is the probability for each site to undergo a substitution during a given spike.
- Additionally, to test the robustness of the method in the face of model misspecification, we have simulated the evolution of sequences under a model where the clock is relaxed, that is, varies in each branch by a factor *e* which is drawn uniformly in [−*ϵ, ϵ*].

We have proposed one possible inference method to check in which circumstances the signal coming from spikes could be distinguished from the signal coming from gradual molecular evolution, when the clock is strict and also when it is relaxed.

The results of the inference on simulated datasets show that the method is able to perfectly identify spikes as soon as *κ* is above a certain threshold 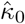 and *ϵ* is below a certain threshold 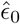. Decreasing *κ* below its threshold makes the effect of spikes indistinguishable from the inherent noise of the (even strict) clock, which results in false negatives (true spikes not detected). Increasing *ϵ* above its threshold makes the variations of the clock indistinguishable from spikes, which results in a number of false positives increasing with *ϵ* (false spikes inferred).

Specifically, for a fixed value 0.08 (per bp per My) of the substitution rate *µ*, the spikes in a 2 kb long sequence on a phylogeny with 32 tips and crown age 8.78 My are perfectly inferred (no false negative, no false positive) as soon as 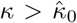, for a clearcut threshold 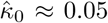 when the clock is strict (*ϵ* = 0). The results also show that for *κ* of the order of 0.03 − 0.05, no false negative is expected when *ϵ* is below the threshold 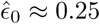, that is 25% variation of the molecular clock in each branch.

The empirical values 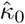 and 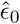 of thresholds are not informative *per se* since they depend on the tree and on the values of other parameters, including molecular clock and sequence length, but can be compared to the theoretical threshold values *κ*_0_ and *ϵ*_0_ predicted in Section 2.5, Equations (2) and (3). Using the parameter values used for the simulations *N* = 2, 000 bp, *µ* = 0.08 per bp per My and *L* = 3 My (average depth of spikes in the simulations) predicts the following threshold values (taking *κ* = 0.04 in the computation of *ϵ* _0_)

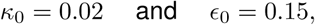

which have the same order of magnitude as the empirical values 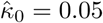 (actual inference is a little worse than predicted) and 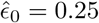 (actual inference is a little better than predicted).

In the snake dataset studied, the sequence length and the substitution rate estimated are equal to *N* = 477 and *µ* = 2.73 10^−3^ per My for CRISP and *N* = 348 and *µ* = 2.04 10^−3^ per My for SVSP. For a typical branch length of this phylogeny, say *L* = 10 My, these values correspond to the same detection threshold of spike amplitudes for the two loci equal to *κ*_0_ = 7.6 10^−3^. The values of *κ* estimated from the alignment are well above this threshold, ranging from 0.035 to 0.065, indicating that the signal of spikes was strong enough to be contrasted from clock-like evolution.

Furthermore, it should be useful to check whether the range of parameter values for which spikes are detectable is in agreement with typical empirical values of the molecular clock and of its temporal variance. One difficulty is that the variance of the molecular clock measured in empirical studies is potentially altered by the actual presence of spikes. One exception is the study by Lartillot *et al.* (2016) on mixed clocks, where clocks are modelled by a stochastic process combining:

- A correlated part embodied by a geometric Brownian motion with infinitesimal variance *σ*^2^ and
- A non-autocorrelated part, where the clock is accelerated or decelerated by an independent, random factor on some edges of the phylogeny.

Interestingly, the latter process is formally similar to the effect of spikes so that the measure of *σ*^2^ (temporal variance of the molecular clock in the former process) made by this method should be immune to the presence of spikes. The typical value of *σ*^2^ estimated by Lartillot *et al.* (2016) on a phylogeny of 105 placental mammals is 1.5 per 100 My (exact average value provided by personal communication of N. Lartillot). In Section 2.5, Equation (4) gives a criterion for distinguishing the effect of spikes from that of a Brownian-like clock. If we replace by their standard values the two parameters in the right-hand-side of this inequality, namely *µ* = 10^−2^ per My and as seen previously *σ*^2^ = 1.5 10^−2^ per My, it becomes

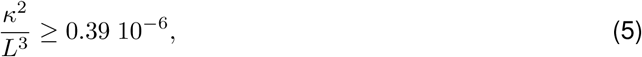

where *L* is measured in My. The inequality (5) constrains the amplitude *κ* of spikes that can be distinguished from clock uncertainty given the branch length *L*. The smaller *L* and the more easily spikes can be detected on that edge. For *L* = 1 My, the minimal *κ* is 6 10^−4^ and for *L* = 10 My, the minimal *κ* is 0.02.

Sticking to these values of *µ* and *L*, and for a sequence of *N* = 1 kb, the minimal amplitude *κ*_0_ of spikes that can be distinguished from the strict clock-like molecular evolution is *κ*_0_ = 6 10^−3^ for *L* = 1 My and *κ*_0_ = 0.02 for *L* = 10 My. These figures show that for sequences at least 1 kb long, whenever spikes can be distinguished from strict clock-like evolution (*κ* ≥ *κ*_0_), they can also be distinguished from the effects of temporal variations of the clock.

### 4.2 Distribution of spikes along the genome: Insight into speciation modes

Since the seminal work of Zuckerkandl and Pauling (1962), standard models of molecular evolution posit that substitutions accumulate through time at a constant rate known as the molecular clock. To account for empirically observed departures from strict clock-like molecular evolution, it is common to assume that the clock itself can vary through time (relaxed clock).

In this paper, we have drawn inspiration from the punctuated equilibrium theory of trait evolution to propose an alternative way of accounting for non-clock-like molecular evolution. We have considered two unconventional hypotheses:

1. Sporadically across the phylogeny, periods of fast accumulation of substitutions can occur, seen as instantaneous events at the macro-evolutionary scale called spikes;
2. Spikes co-occur with the speciation process.

In contrast with relaxed clock models which rely on *ad hoc* assumptions agnostic to evolutionary processes, the spike model of molecular evolution is based on two bottom-up descriptions of molecular evolution in relation to speciation: (i) ecological speciation with gene flow can result in the rapid formation of genomic islands of differentiation; (ii) hybridization with a distant lineage can result in the adaptive introgression of ancient genetic variation into the focal lineage.

Additionally, a subsequent distinctive feature of the spike model is that it draws information not only from the sequence alignment but also from the diversification process itself. Because spikes occur at speciation, clades which are more speciose are expected to be more prone to spikes.

The two evolutionary processes putatively causing spikes can hypothetically be identified via two distinctive genome-wide signatures. In the case of ecological speciation, substitutions due to the spike are localized in specific genomic regions (genomic islands) which can be detected for example by scanning the genome in search for *F*_*st*_ outliers (Seehausen *et al.*, 2014). In the case of distant hybridization, the amplitude of the spike is expected to be approximately uniform among introgressed loci, equivalent to the effect of the gradual accumulation of substitutions during twice the divergence time between the donor and receiver lineages, which can also be assessed by genome scans (Osada and Wu, 2005). In the application of our method to venom proteins given here, only a couple of loci have been analyzed, preventing us from diagnosing the specific causes of the spikes inferred via genome-wide signatures.

In future work, the search for empirical evidence of spikes will include: (i) measuring the rate of differentiation in typical genomic islands of speciation and (ii) measuring genetic distance between a typical allele introgressed from a distant lineage and the removed allele. These studies would give empirical estimates for the value of *κ*.

### 4.3 Evolution of snake venom proteins

Venom is a key adaptation for many snakes that facilitates the capture and the predigestion of prey. While the majority of the well-known venomous snakes are part of the Viperidae and Elapidae families, venoms seem to have originated earlier in squamate evolution, around 170 million years ago (Fry *et al.*, 2012). Thus, snakes together with Anguimorpha (*i.e.*, monitor lizards and alligator lizards) and Iguania (*i.e.*, iguanas, chameleons and agamid lizards) constitute the venom clade (Toxicofera) (Fig. 5). According to this hypothesis, all members of this clade may be venomous to a certain degree. Non-venomous snakes still possess venom proteins, but lack a delivery method or harmful venom (Fry *et al.*, 2012). Venom proteins evolved throughout the Toxicofera with a lot of evidence supporting positive selection (Župunski and Kordiš, 2016), accelerated evolution (Siigur *et al.*, 2001), shifts in the splicing site or in the reading frame due to small deletion or insertion (Doley *et al.*, 2009) and large effect mutations (Vaiyapuri *et al.*, 2011).

Venoms are composed of several proteins able to attack biological pathways. A very recent study investigated the evolution of toxin combinations among snakes (Barua and Mikheyev, 2019). Despite some evidence supporting the convergent evolution of envenomation strategies that suggests the influence of ecological filtering, the variation in toxin component (transcriptome) was clustered into three distinct adaptive optima and showed a clear association with phylogeny. These adaptive optima represent three distinct envenomation strategies. The Elapids’ venoms were dominated by TFTx (three-finger toxin) proteins which damage the nervous system (Kini and Doley, 2010). Vipers’ venoms were mainly dominated by either SVMP (snake venom mellanoprotease) proteins which cause hemorrhage and ischemia (Urs *et al.*, 2014) or the combination of SVSP (snake venom serine protease) and PLA2 (phospholipase A2) proteins which disrupt the haemostasis and promote cell lysis respectively (Meier and Stocker, 1991; Nicolas *et al.*, 1997). Colubrids’ venoms appeared to be the most diverse in composition, employing all of the three different strategies.

While showing no spike for R35 (supplementary Fig. S1), suggesting a clock-like evolution of this gene used in molecular reconstruction, our model of evolution with spikes highlights previously described envenomation strategies by showing differential spiking patterns on toxin evolution among the Toxicofera (Fig. 5) reflecting differential selective pressures. The inferred spiking patterns suggest stronger selective pressure on the neurotoxin (CRISP) among Elapids and NFFC (non-front fanged colubrids) and on the hemotoxin (SVSP) among Viperids. These results are in line with the study described above which examined venom transcriptomes (Barua and Mikheyev, 2019).

### 4.4 Development opportunities

The current implementation of the MCMC is only efficient enough to perform the exploratory tests of the method that we presented in this paper. In its current state, it already takes about 1 or 2 days to complete a 10^6^-step chain on a 50-leaf tree displaying a 2 kb alignment. This performance is not compatible with genome-wide analyses that one should like to do in the future. First, spiking signal from multiple loci would help locate spikes on the genome with more confidence. Second, variation of spike density across the genome and spike amplitudes across loci could help disentangle the possible evolutionary causes of spikes (see section 4.2). The performance could be improved at least in three ways. First, there is still room for more carefully designing the MCMC proposal in order to speed up the mixing of the chain. Second, one could use a compiled programming language and rely on already available efficient implementations of likelihood computation for models of molecular evolution (in popular softwares such as BEAST or RevBayes). Third, a genome-wide analysis can be done in two steps. A first step would consist in rapidly locating spikes affecting multiple loci thanks to a distance-based method, similarly as done by Tamura *et al.* (2012). The second step would be locus-specific and use an improved version of the MCMC presented here.

### 4.5 Conclusion

In recent years, a popular approach for uncovering the process of species diversification has relied on lineage-based models where species are seen as particles that can give birth (speciation) or die (extinction) at rates possibly depending on time (Morlon *et al.*, 2011; Stadler, 2011), species age (Lambert and Stadler, 2013; Alexander *et al.*, 2015), speciation stage (Etienne *et al.*, 2014; Lambert *et al.*, 2015), clade and number of co-occurring species within clade (Etienne *et al.*, 2011; Rabosky *et al.*, 2014), a known (Maddison *et al.*, 2007) or unknown (Beaulieu and O’Meara, 2016) trait. The methods associated with these models have been widely applied to large scale phylogenies to infer past diversification, to estimate diversification rates and to characterize how these rates depend on the aforementioned variables.

On the other hand, the novel field of speciation genomics has made progresses on our understanding of the trace left by the speciation process on genomes and conversely on the inference of modes of speciation from genomic data (Feder *et al.*, 2013; Seehausen *et al.*, 2014; Roux *et al.*, 2016; Meier *et al.*, 2017b).

These two classes of methods have specific benefits and shortcomings. The first class of methods can handle the information on phylogenetic relationships between many species but relies on the knowledge of a single phylogeny, which can result in problems of identifiability and false positive associations between traits and rates (Rabosky and Goldberg, 2015; Moore *et al.*, 2016; Louca and Pennell, 2019). The second class of methods can handle the information coming from whole-genome sequences but usually only restricted to a handful of species. We believe that modern approaches will combine these two classes of methods thanks to models coupling the processes of diversification and of molecular evolution and to methods drawing on both large-scale phylogenies and on the rich signal contained in their genomes. We hope that the present work will pave the way for such approaches (see also Tank *et al.*, 2015; Marin *et al.*, 2019).

We have argued here that two specific modes of speciation should leave a characteristic signature on genomes: ecological speciation with gene flow (localized spikes with varying amplitudes) and hybridization with a distant lineage (sparsely distributed spikes of uniform amplitude). To use directly sequences as witnesses of the diversification process, we will need in the future additional general hypotheses of this sort and scalable inference methods capable of testing them.

## 5 Material & Methods

### 5.1 Inference on simulated data

We performed tests of the inference method on simulated datasets under known parameter values, and assessed our ability to retrieve those parameters using our inference protocol. Because the parameter space is huge and each run of the MCMC takes time, we restricted ourselves to manually chosen parameter values, and varied one parameter at a time (Table 1). For each parameter combination, (*λ, d, ν, α, β, κ*), we used our code to simulate one dataset (a reconstructed spiked tree, and an alignment evolving on it), and estimated the marginal posterior distribution of the parameter under scrutiny. We fixed *f* = 1 in all these analyses. We fixed the time of origin 10 units of time in the past, and simulated sequences of length 2 kb. On each dataset, the MCMC is run for 10^6^ generations, and a burn-in of 3 10^5^ generations is discarded to estimate the posterior probability. Results are reported in Fig. 3.

We performed another set of analyses in order to assess when the amplitude *κ* of spikes is large enough for the spikes to be detected. For these analyses we varied *κ* while holding other parameters constant. These analyses are described below, corresponding to the case when *ϵ* = 0, and the results are reported in Fig. 3.

### 5.2 Robustness to model misspecification

We tested how heterotachy, *i.e.*, variation in lineage substitution rates across lineages, affects our inference method. We model this variation with a one-parameter relaxed clock, where each branch length *L* of the reconstructed tree is replaced by *L*(1 + *e*) where *e* is sampled independently in each branch from the uniform distribution on [−*ϵ*, +*ϵ*].

The robustness to model misspecification is expected to depend both on the level of heterotachy (value of *ϵ*) but also on other model parameters. For simplicity, we only varied the parameter *ϵ* ∈ {0.0, 0.1, 0.2, 0.3, 0.4} and the amplitude of spikes *κ* ∈ {0.03, 0.04, …, 0.13}. We first simulated a spiked tree with *λ* = 1, *d* = 0.8, *f* = 1 and *ν* = 0.1. We fixed this reference tree, harboring 32 tips and 4 spikes. For each parameter combination (*ϵ, κ*), we simulated corresponding alignments of 2 kb with *α* = 0.02 and *β* = 0.03.

Next, we tested the ability of the MCMC algorithm to sample from the posterior distribution, knowing the alignment at the tips of the tree and the tree for each parameter combination. We checked for convergence (ESS scores), and averaged and rounded the number of spikes at each node over the whole distribution after a burn-in of 1,000 generations (out of 50,000 generations).

### 5.3 Snake venom proteins evolution

We evaluated the spiking pattern of two proteins, SVSP (snake venom serine protease) and CRISP (cystein-rich secretory protein), involved in different envenomation strategies among the Toxicofera (Barua and Mikheyev, 2019). We retrieved the alignments of SVSP and CRISP for 46 and 53 species of snakes respectively from Perry *et al.* (2018). These two genes were available in a single copy (orthologous sequences) for most of the species (80%) (see Perry *et al.*, 2018). When there were several copies, we selected orthologous sequences using a phylogenetic criterion. Sequences correctly located in their family clade according to a maximum likelihood analysis (Perry *et al.*, 2018) were kept and phylogenetically misplaced sequences were discarded. We thus obtained a single copy of each gene for each species.

Gaps and codons containing undefined bases were removed from the alignments. After correcting for synonyms (supplementary table S1), calibrated trees were downloaded from the timetree database (http://www.timetree.org/) (Kumar *et al.*, 2017). We assessed the sampling fraction of each family, using taxonomic information from the Reptile Database (Uetz *et al.*, 2019), from which we deduced the global sampling fraction used in our analyses (parameter *f*). The final data set (alignments and corresponding timetrees) comprised 46 sequences of 348 bp (of which 286 sites were variable) for SVSP and 53 sequences of 477 bp (of which 347 sites were variable) for CRISP. Additionally, we downloaded 37 R35 (Orphan G protein-coupled receptor R35) sequences of 519 bp (of which 281 sites were variable) from GenBank (supplementary table S3). The R35 sequence alignment was performed with ClustalW2 (Larkin *et al.*, 2007) implemented in BioEdit (Hall *et al.*, 2011) and then manually refined.

We used our model of molecular evolution with spikes to infer the spiking pattern of each gene (CRISP, SVSP and R35). Because we studied coding sequences, we added the possibility to infer the molecular evolution parameters (*α, β* and *κ*) independently at each codon position in our code (Supplementary material). To reach convergence we performed and combined 4 runs of 10,000,000 generations of the MCMC algorithm (ESS scores > 200, burn-in = 10,000).

## ACKNOWLEDGMENTS

The authors are grateful to Todd A. Castoe for providing them with the venom gene alignments. They thank Nicolas Lartillot, Tanja Stadler, Guillaume Achaz, Marie Manceau, Grégory Nuel, Ana C. Afonso Silva, Leandro Aristide, Laure Ségurel, Richard Durbin and Nicolas Vidal for fruitful discussions on the topic of the paper. AL, MM and JM thank the Center for Interdisciplinary Research in Biology (CIRB, Collège de France) for funding. AL, HM and JM thank the LabEx MemoLife for funding (project “Genomics of species diversification”).

## Supplementary Material

### A MCMC Implementation

We describe here the design of the initialization of the chain, and the movement proposal, in the MCMC aimed at sampling our posterior probability.

#### A.1 Initialization of the chain

Without prior knowledge on parameter values, we initialize them by drawing from their prior distributions. The initialization of the spike configuration is done more carefully, by annotating branches on which there are more substitutions than expected in order to tune an appropriate initial distribution *g*.

We call 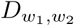 the random variable counting the number of sites that are in a different state at two extremities *w*_1_ and *w*_2_ of a branch. More precisely, we wish to compare

1. The expected number of differences on the branch, 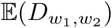.
2. The expected number of differences conditional on present-day data, 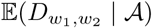.

The first quantity is derived analytically from the knowledge of the substitution model, while the second one is numerically computed through two depth-first traversals of the tree, as outlined in Friedman et al. (2002). In a first postorder traversal, we compute the probability of nucleotide states at the two ends of each branch, conditioned on the states of the tips subtended by that branch. In a second preorder traversal, we compute the probability of nucleotide states at the two ends of each branch, conditioned on the states of all the tips of the tree. Summing over the sequence the probabilities that nucleotides are different at the two ends of a branch gives 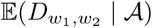.

We measure the difference between the two quantities, informing us on the departure from what we would expect in the absence of spike

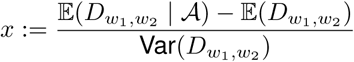

When *x* is large, we want to place a spike with higher probability along the branch. We chose to place a spike on a branch with a probability *l*(*x*), where *l* is the logistic function

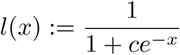

We further chose *c* such that, as soon as *x* is large enough (*e.g.*, we took *x* > 1.96, but any value of the same magnitude could be chosen), there is a probability *l*(*x*) > *ν* to place a spike. This gives us,

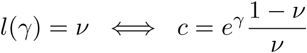

Because in the model spikes can occur at a node of the reconstructed phylogeny and also inside branches (at ‘hidden’ nodes), we would like *g* to have the same behavior. We assume that *g* is the convolution of a Bernoulli distribution with parameter *l*(*x*) and a Poisson distribution with parameter *ζ* defined in main text, Equation (1).

#### A.2 Movement proposal

We now describe the movement proposal, aimed at drawing a new state (𝒮′, *λ*′, *d*′, *ν*′, *κ*′, *α*′, *β*′) from a previous state (𝒮, *λ, d, ν, κ, α, β*) at each step of the chain.

First, we chose not to change the full state at each step. We rather change only the following subsets with probabilities 1/8: (*λ, d*), (*ν, κ*), (*α, β*), (*λ, d, ν, κ*), (*λ, d, α, β*), (*ν, κ, α, β*), (*λ, d, ν, κ, α, β*), or 𝒮.

When they are changed, each of the 6 real parameters *λ*′, *d*′, *ν*′, *κ*′, *α*′, *β*′ will be drawn in a Gaussian distribution centered respectively on *λ, d, ν, κ, α, β*, with a specific variance and conditioned on staying in a specific interval.

- Parameters *λ, d* are conditioned on staying in (0, 5) and their Gaussian has variance 0.5.
- Parameters *ν, κ* are conditioned on staying in (0, 1) and their Gaussian has variance 0.1.
- Parameters *α, β* are conditioned on staying in (0, 0.1) and their Gaussian has variance 0.01.

Last, the following law governs the transition from the spike configuration 𝒮 to another configuration 𝒮′,

1. If 𝒮 has *n*_*s*_ > 0 spikes, then:
  a. With probability 0.05, a number *U* of spikes are deleted chosen uniformly among all spikes, where *U* is uniform in (1, *n*_*s*_).
  b. With probability 0.95 a number 1 + *P* of spikes is added to the tree, uniformly among all nodes, where *P* is a Poisson random variable with parameter 1.
2. Otherwise, only the previously described addition of spikes is performed.

This ends the description of the movement proposal, which determines the mixing efficiency of the MCMC. Note that our operator depends on parameters that can be chosen by hand so as to achieve a faster convergence: (i) the probabilities to change each parameter can be adjusted so as to ensure that each parameter on average moves as often as others, (ii) variances of the Gaussian distributions adjust the size of the steps for the new parameter set, and (iii) the parameter of the Poisson distribution as well as the propensity to add or remove spikes can also be tuned.

#### A.3 Structure of the code

The code used in this study is freely available in the GitLab repository https://gitlab.com/MMarc/spike-based-clock/. It is written in Python and organized in the following core / accessory / test files.

Core files are:

**nt.py** contains all functions handling nucleotides, sequences, and alignments.

**evolmol.py** contains the description of the model of molecular evolution used here, namely, K80.

**evoltree.py** contains all necessary functions to simulate molecular evolution along a tree, or compute the probability density of an alignment having evolved along a tree.

**tree.py** contains functions simulating birth-death trees and computing the probability density of birth-death trees.

**spike.py** contains functions for simulating spikes along a tree, as well as moving spikes according to different schemes during the MCMC implementation.

**mcmc.py** is probably the most important file, or at least the one that people might be the most interested in modifying. It contains two MCMC implementations: the first one considering that all nucleotides along the sequence evolve identically, and the second one considering that the first, second and third position of a codon evolve with different parameters. The initialization, moves, and priors could be changed according to the problem under study.

Accessory files are:

**graphics.py** contains a few graphic functions to display spikes along trees, and to graphically analyze the MCMC outputs.

**export.py** which allows one to average spikes at each node, over the posterior.

Test files are:

**try_inferencesOnSimulations.py** to run a MCMC on a simulated dataset.

**CRISP_ordered.fasta** the alignment of CRISP sequences.

**try_CRISP_3pos.py** to run a MCMC on the CRISP dataset.

### B Quantifying the ability to detect spikes

In this section we aim to give a rule of thumb on the model parameters to indicate when the effect of spikes can be distinguished from the stochasticity of clock-like substitutions and from temporal variations of the molecular clock.

Let us focus on a single branch of length *L* and assume as in the main text that the molecular clock *M* on this branch is equal to *µ* = *α* + 2*β* plus some random variation *eµ*, where *e* is uniformly distributed in [−*ϵ*, +*ϵ*] (which is equivalent to assuming uncertainty on branch length, as done in Material & Methods). If we assume the presence of one single spike on this branch (either at the mother node or inside the branch), then the number of mutations accumulated on the branch is the sum of *S* and *C*, where *S* is the number of mutations due to the spike and *C* the number of mutations due to the clock. If *N* denotes the total number of target sites, then *S* is binomial with parameters *N* and *κ* and *C* is Poisson with parameters *MNL* (conditional on *M*, which is itself random). The stochastic effects of *S* and *C* can be discriminated if *S* + *C* is statistically different of *C*, that is, if the mean *κN* of *S* is large compared to both the standard deviations of *S* and *C*, hereafter denoted *σ*_*S*_ and *σ*_*C*_ respectively. More specifically, since *S* and *C* are Binomial and Poisson variables respectively, a CLT (central limit theorem) approximation applies and our criteria for identifiability of spike vs clock effects read

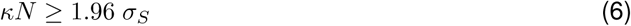

and

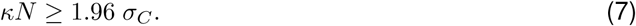

Standard computations yield

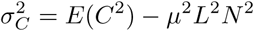

and because conditional on *M, C* is Poisson with parameter *MLN*,

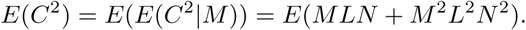

Now *E*(*M*) = *µ* and *E*(*M* ^2^) = *µ*^2^ + *ϵ*^2^*µ*^2^/3, so we get

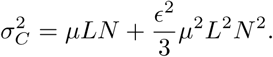

On the other hand,

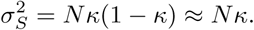

So Criterion (6) is equivalent to having *κN* ≥ 1.96 *σ*_*S*_, which is equivalent to *κN* ≥ (1.96)^2^, that is *κN* ≥ 3.84. On the other hand, Criterion (7) reads

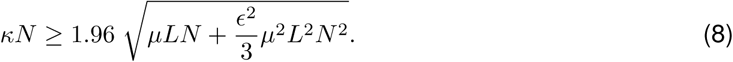

In effect, one must at least have 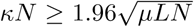 to enforce statistical inference of spikes when the clock is strict (*ϵ* = 0). This last inequality reads *κ* ≥ *κ*_0_, where

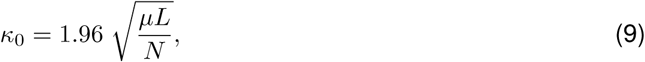

When *ϵ* ≠ 0, (8) implies to have at least 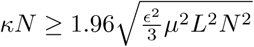, which reads *ϵ* ≤ *ϵ*_0_, where

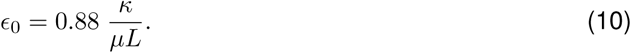

In a geometric Brownian motion model of relaxed clock, the substitution rate varies through time like 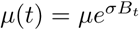, where *B* is a standard Brownian motion. Then for small *L*, writing *µ*(*L*) ≈ *µ*(1 + *σB*_*L*_), the variance of *C* becomes

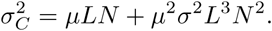

The same reasoning as previously leads to the following criterion for *σ* required for variations of the molecular clock to not blurr the spike signal:

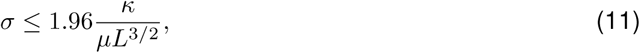

or equivalently

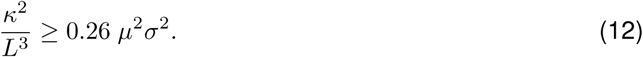

